# Population reweighting and reporter distortion obscure ribosome regulation during translation inhibition

**DOI:** 10.64898/2026.09.30.755718

**Authors:** Ruizhe Li, Kieran Abbott, Somenath Bakshi

## Abstract

Bacteria dynamically adjust their ribosome allocation as growth conditions change, but how this regulation unfolds within individual cells remains poorly understood. Established relationships between ribosome allocation and growth are largely derived from population measurements under steady-state conditions, which may not capture the trajectories followed by individual cells during perturbation. Differential growth can reweight heterogeneous cellular states over time, causing population-level measurements to diverge from the regulatory trajectories followed by individual cells. Here, we use lineage-resolved single-cell microscopy to follow growth and ribosomal reporter dynamics in *Escherichia coli* during translation inhibition without growth-dependent reweighting of the sampled lineages. Unexpectedly, fluorescent protein fusions to the ribosomal proteins S2 and L9 decrease in fluorescence following chloramphenicol treatment, whereas transcriptional reporters driven by the ribosomal promoters initially increase. The decrease in ribosomal fusion fluorescence persists across multiple fluorescent proteins and is reproduced during tetracycline treatment, indicating that it is not specific to a particular fluorophore, ribosomal protein or translation inhibitor. Although some fusion reporters retain a relationship with cellular growth within physiological states, this relationship is itself reporter-dependent. These results reveal two independent requirements for quantitative measurements of dynamic cellular responses: longitudinal sampling is needed to distinguish within-cell regulation from growth-dependent population reweighting, and reporter behaviour must be validated when the perturbation alters the processes generating the measured signal.

## Introduction

Ribosome allocation is a central determinant of bacterial growth and physiological adaptation^1^. During balanced growth, the fraction of cellular resources allocated to ribosomes varies systematically with growth conditions, giving rise to well-established quantitative relationships between ribosome abundance and growth rate^1–3^. Translation inhibition produces a strikingly different relationship: as translational activity is progressively inhibited, cells increase their allocation to the ribosomal sector even as growth slows^1,4^. These relationships have provided a quantitative framework for understanding bacterial resource allocation and have motivated the use of fluorescent reporters of ribosome abundance and organisation within cells^5,6^, opening the possibility of following ribosome allocation as cells adapt to changing environments.

These growth laws describe relationships between population averages across steady-state conditions and do not, by themselves, establish how ribosome allocation relates to growth among individual cells within one condition. Recent single-cell measurements have indeed revealed substantial variability in ribosome allocation and only weak correlations between ribosome concentration and growth rate within a given condition^7^. In resource-allocation frameworks, growth depends on balancing investment in ribosomal and metabolic sectors, but the relationship between proteome allocation and growth is not universal and can depend on the underlying regulatory architecture^8^. Applying population-level relationships to individual cells, particularly during a dynamic perturbation, therefore requires direct experimental assessment.

Extending these relationships from balanced growth to dynamic perturbations presents a distinct measurement problem. Individual cells can enter a perturbation in different physiological states and subsequently differ in their growth, arrest and adaptation trajectories. Because faster-growing cells progressively contribute more descendants to a population, these differences alter the composition of the population over the course of a measurement. A sequence of population-averaged measurements can therefore reflect both regulation occurring within cells and growth-dependent changes in population composition, and need not represent the regulatory trajectory followed by any individual cell or lineage. This is particularly relevant during translation inhibition, where ribosome allocation, translational activity and growth are strongly coupled and can change substantially over the course of the response.

Resolving ribosome regulation dynamically therefore requires three experimental capabilities. First, individual cellular lineages must be followed continuously so that within-cell regulatory changes can be distinguished from changes in population composition. Second, those lineages must experience the same temporal profile of environmental perturbation, allowing differences in their responses to be attributed to cellular physiology rather than variation in exposure. Third, ribosome allocation must be quantified continuously within the same cells as their physiological state changes. Longitudinal single-cell measurements using single-cell microfluidics combined with time-lapse microscopy provide a framework for satisfying the first two requirements. An adapted mother-machine microfluidics design allows the same bacterial lineages to be followed continuously while the extracellular environment is rapidly changed, enabling reporter dynamics and cellular physiology to be measured through a perturbation without growth-dependent reweighting of the sampled population^9^. Fluorescent fusions to ribosomal proteins provide an attractive approach to the third requirement by enabling ribosome-associated signals to be followed continuously in living cells.

Here, we combine lineage-resolved microfluidic measurements with fluorescent reporters of ribosome transcription and translation to examine dynamic ribosome allocation during translation inhibition in *Escherichia coli*. We find that fluorescent ribosomal protein fusions exhibit unexpected dynamics during antibiotic treatment, limiting their interpretation as quantitative reporters of dynamic ribosome allocation. We test the generality of this discrepancy across reporter designs and translation inhibitors, and compare these dynamics with transcriptional regulation reported from a ribosomal promoter. Translation inhibition represents a particularly stringent test of fluorescent reporters because the perturbation directly affects the protein-synthesis machinery required to generate the reporter signal. In a companion study, Zodage et al. independently investigate the basis of this discrepancy using biochemical measurements and an orthogonal fluorescence-LacZ reporter, showing that fluorescence output can become uncoupled from reporter abundance during translation inhibition^10^. Together, the two studies reveal complementary aspects of the challenge of quantitatively interpreting fluorescent ribosome reporters when the perturbation directly targets the processes required to generate their signal.

## Results

Ribosome allocation occurs within individual cells in response to their physiological state, but population-level measurements do not necessarily report the regulatory trajectory followed by individual cells. When cells differ in their physiological states and growth responses, faster-growing cells progressively contribute more descendants to the population, causing their states to become increasingly represented in population-level measurements (Fig. 1a). As a result, even when a population and a set of tracked lineages originate from the same initial distribution of cellular states, their average trajectories can diverge over time. We therefore used a simple illustrative model to examine how differential growth alone can separate population-level measurements from the equally weighted trajectories of the initially sampled lineages.

**Figure 1:**
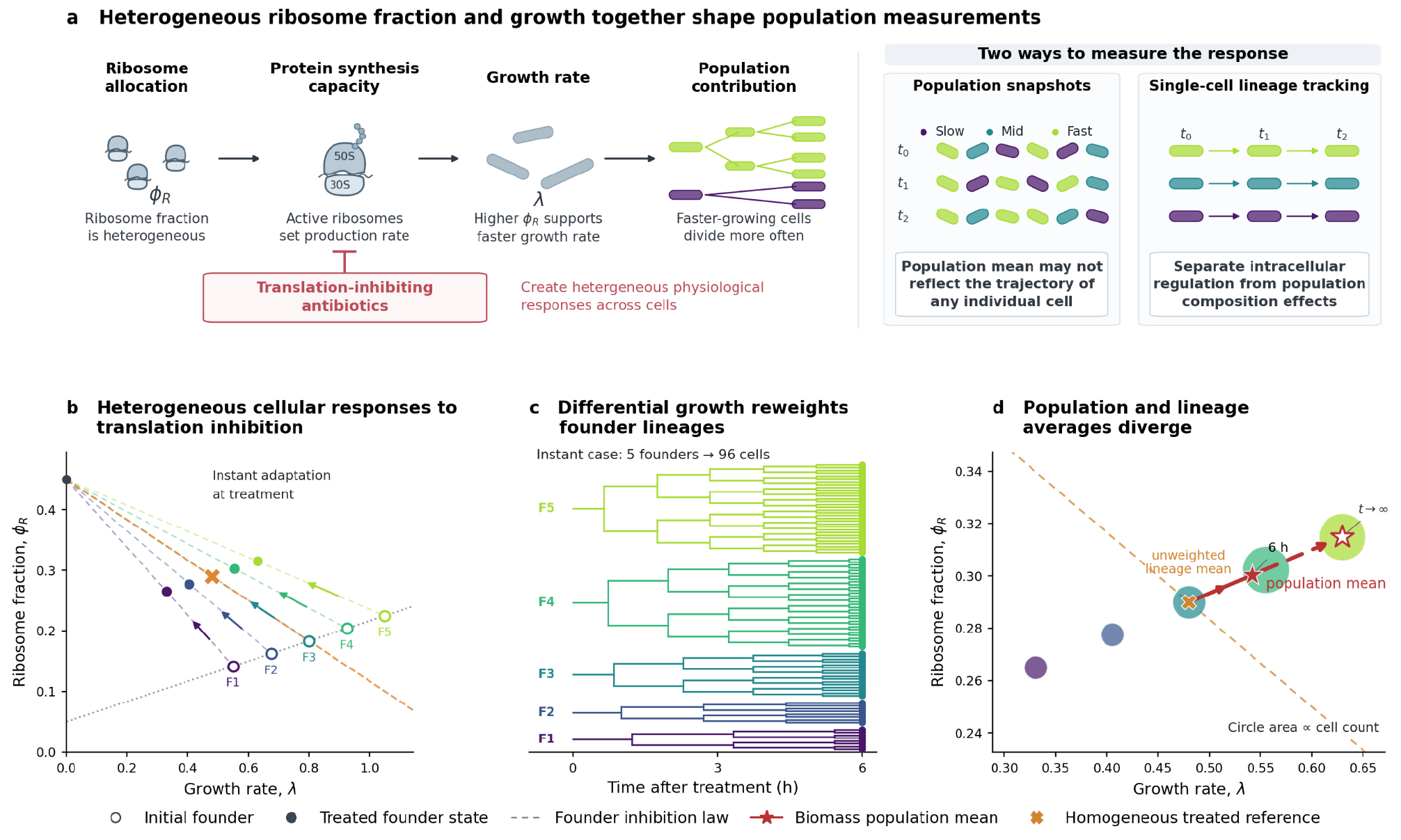
Differential growth can separate population-level measurements from the trajectories of individual lineages. **a)** Conceptual distinction between population snapshots and longitudinal single-cell lineage tracking. When cellular physiology and growth are heterogeneous, faster-growing lineages contribute progressively more descendants to an evolving population, whereas lineage tracking follows the initially sampled cells with equal weight. **b)** Illustrative model of heterogeneous founder cells undergoing translation inhibition. For simplicity, cellular states before and after treatment are assigned using the population-level *E. coli* relationships between growth rate (*λ*) and ribosome fraction (Φ_*R*_). This provides defined initial and treatment-associated states for each lineage but is not intended to imply that the population-level growth law constitutes a deterministic single-cell relationship between *λ* and Φ_*R*_. Open circles indicate the initial states of founder cells and filled circles their treated states; coloured trajectories connect the corresponding cellular states. Dashed lines indicate the growth-law relationships before and during translation inhibition, and dotted lines indicate the mean reference states. **c)** Expansion of the founder lineages over 6 h following treatment. Differences in growth history generate different numbers of descendants from initially equally represented founder cells, progressively reweighting their contributions to a population-level measurement. **d)** Cellular states after 6 h, with circle area proportional to the number of descendants arising from each founder lineage. Under this illustrative parameterisation, faster-growing lineages become over-represented, causing the growth-weighted population mean to diverge from the equally weighted mean of the original founder lineages.

To illustrate this effect, we considered a simple model in which individual founder cells occupy heterogeneous physiological states characterised by a ribosome fraction, Φ_*R,i*_ (*t*), and an instantaneous growth rate, *λ*_*i*_(*t*) (Fig. 1b). For simplicity, the initial and treatment-associated states of each cell were assigned using the population-level E. coli relationships between growth rate and ribosome allocation before and during translation inhibition^1,11^. This assumption provides a convenient way to define where each lineage starts and the state towards which it adapts, but is not intended to imply that the population-level growth law constitutes a deterministic single-cell relationship between Φ_*R*_ and *λ*. Starting from these heterogeneous initial states, each founder lineage follows its own physiological trajectory during translation inhibition, while the biomass arising from lineage *i* evolves according to 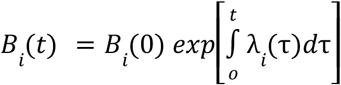.

A population-level measurement of ribosome allocation therefore corresponds to a growth-weighted average,

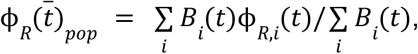

whereas a lineage-resolved measurement gives each lineage equal weight,

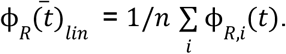

These quantities are identical only when the lineages contribute equally to the population, or when their physiological states are unrelated to their cumulative growth. In the model, differential growth progressively reweights the initially sampled states, enriching the population for lineages that accumulate more descendants (Fig. 1c). Consequently, the population-weighted mean diverges from the equally weighted lineage mean, despite both measurements originating from the same initial distribution of cellular states (Fig. 1d). Under the illustrative parameterisation used here, this reweighting shifts the population mean towards the faster-growing states.

Because the steady-state growth-law relationships do not specify how individual cells move between their initial and treatment-associated states, we next asked whether the population-lineage discrepancy depended on the assumed adaptation trajectory. We therefore considered two limiting scenarios in which growth rate responds before ribosome allocation, or ribosome allocation responds before growth rate (Fig. SI1a,b). These assumptions generate different paths through physiological state space and consequently different transient population-average trajectories. However, in both cases differential growth progressively reweights the contribution of the founder lineages, causing the population average to diverge from the equally weighted lineage average (Fig. SI1c). Thus, while the precise transient trajectory depends on how growth and ribosome allocation adapt to treatment, the central sampling effect does not: <u>population-level measurements combine within-lineage physiological change with growth-dependent changes in lineage representation.</u>

Model parameters are provided in table 1.

**Table 1.** List of parameter values used in agent-based simulation of growth-dependent population reweighting.

| Parameter | Symbol | Value |
| --- | --- | --- |
| Number of founder cells | $n$ | 5 |
| Mean initial growth rate | $\mu_\lambda$ | $0.8 \text{ h}^{-1}$ |
| Initial growth rate standard deviation | $\sigma_\lambda$ | $0.18 \text{ h}^{-1}$ |
| Baseline ribosome fraction | $\phi_{base}$ | 0.05 |
| Growth-law coefficient | $\kappa_t$ | $6 \text{ h}^{-1}$ |
| Maximum ribosome fraction | $\phi_{max}$ | 0.45 |
| Residual growth factor | $q$ | 0.6 |
| Trajectory sampling interval | $\Delta t_{sampling}$ | $0.02 \text{ h}$ |
| Adder constant | $\Delta$ | $2 \text{ a. u.}$ |

**Table 2.** List of strains constructed for this study.

| Strain name | Gene | C-terminally fused FP | Background strain | Resistance gene |
| --- | --- | --- | --- | --- |
| RF4 | *rpsB* (S2) | SCFP3A | SB8 | kanR, FRT-flanked |
| RF5 | *rpsB* (S2) | mVenus-NB | SB8 | kanR, FRT-flanked |
| RF6 | *rpsB* (S2) | mScarlet-I3 | SB8 | kanR, FRT-flanked |
| RF14 | *rpsB* (S2) | mJuniper | SB8 | ampR, FRT-flanked |
| RF20 | *rplI* (L9) | mJuniper | SB8 | ampR, FRT-flanked |
| RF21 | *rplI* (L9) | mVenus-NB | SB8 | ampR, FRT-flanked |
| RF22 | *rplI* (L9) | mScarlet-I3 | SB8 | ampR, FRT-flanked |

Neither the population nor lineage average is intrinsically the correct description of the system; rather, they answer different biological questions. The population average describes the mean state of the population present at a particular time, after it has been reshaped by differential growth. The lineage average instead describes the mean trajectory followed by the initially sampled cells. <u>Because regulation is enacted within individual cells, understanding how cells regulate ribosome allocation in response to translation inhibition requires the latter: direct measurement of the regulatory trajectories of individual cells</u>. This distinction is particularly relevant during translation inhibition, where ribosome allocation, translational activity and growth are strongly coupled and individual cells can differ substantially in their response to antibiotic exposure. We therefore used longitudinal mother-machine microscopy to follow the same *E. coli* lineages continuously through translation inhibition^9^. This approach maintains individual lineages while allowing rapid changes in the extracellular environment, enabling growth and ribosomal reporter dynamics to be measured along the same cellular trajectories without growth-dependent reweighting of the sampled cells.

Following the same lineages removes growth-dependent changes in sample composition, but it does not by itself guarantee that the measured reporter faithfully represents the underlying biological quantity. We therefore asked whether fluorescent ribosome reporters provide a consistent readout of ribosome regulation when the same cells are followed through translation inhibition. To measure ribosomal regulation along individual cellular trajectories, we followed *E. coli* lineages carrying two complementary reporters during chloramphenicol treatment. The ribosomal protein S2 (RpsB) was C-terminally fused to YFP to provide a ribosome-associated translational fusion signal, while CFP expressed from the *rpsL* promoter reported transcriptional regulation of a ribosomal protein gene (Fig. 2a). Cells were maintained in the mother-machine device and continuously imaged as the medium was switched from balanced growth conditions in M9 medium (supplemented with 0.04% glucose and 0.02% CAA) to sub-MIC chloramphenicol exposure (4 ug/ml). This setup allowed growth and both reporter signals to be quantified in the same lineages throughout the perturbation. Representative timelapse images of the two reporter channels are shown in Fig. 2b, with corresponding single-lineage growth and reporter trajectories in Fig. 2c.

**Figure 2:**
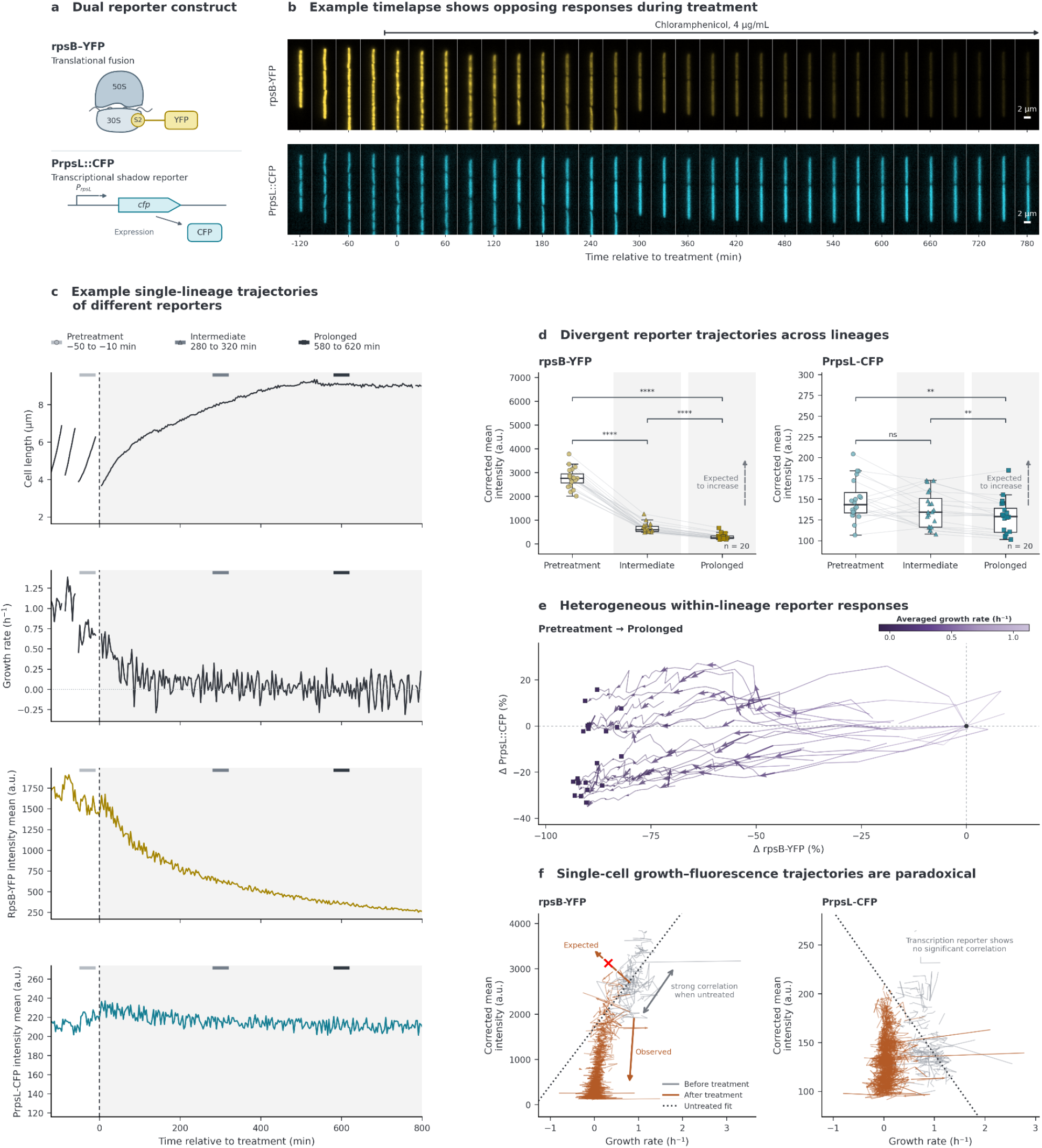
Direct visualisation of the reporter dynamics during translation inhibition. **a)** Schematic of the reporter construct. YFP is fused with the C-terminus of the ribosomal protein S2 (rpsB) and CFP expressed under the ribosomal *rpsL* promoter. **b)** Representative time-lapse fluorescence images of an individual mother-machine lineage carrying the RpsB-YFP translational fusion reporter (top) and the rpsL promoter-driven CFP transcriptional reporter (PrpsL-CFP; bottom) during exposure to chloramphenicol (4 μg ml^−1^). Images are shown at successive time points after treatment addition and illustrate the progressive decrease in RpsB-YFP fluorescence and concurrent increase in PrpsL-CFP fluorescence. **c)** Representative single-lineage trajectory showing cell length, instantaneous growth rate, RpsB-YFP mean fluorescence intensity and PrpsL-CFP mean fluorescence intensity during chloramphenicol treatment (shaded region). Growth rate rapidly decreases following treatment. Over the same cellular trajectory, RpsB-YFP fluorescence progressively declines, whereas the example PrpsL-CFP fluorescence did not change significantly. **d)** Quantification of RpsB-YFP translational reporter (left) and PrpsL-CFP transcriptional reporter (right) fluorescence before treatment (−50 to −10 min), during the intermediate response (280-320 min) to chloramphenicol exposure, and after prolonged treatment (580-620 min). Measurements from the same mother-machine lineages are connected by grey lines. Boxplots show the median and interquartile range and whiskers extend to 1.5× the interquartile range. RpsB-YFP fluorescence decreases progressively during translation inhibition, whereas PrpsL-CFP fluorescence showed only later a significant reduction on average. Adjusted P values for paired comparisons are indicated. **e)** Paired changes in RpsB-YFP and PrpsL-CFP fluorescence within individual lineages during chloramphenicol treatment. Each trajectory represents the change in both reporter signals for a tracked lineage relative to its pretreatment state at the indicated treatment stage and is colourcoded by its spontaneous growth rate averaged across a 20 mins window. 40% lineages show decreased RpsB-YFP together with increased PrpsL-CFP, whereas others show weaker transcriptional induction or decreases in both reporters, revealing heterogeneous within-lineage responses to translation inhibition. **f)** Trajectories of reporter fluorescence against instantaneous growth rate for individual lineages before and during chloramphenicol treatment. Each line represents one tracked lineage; pretreatment trajectories are shown in grey and treatment trajectories in brown. Dotted lines indicate linear fits to the pretreatment measurements (RpsB-YFP r = 0.63, PrpsL-CFP r = -0.53). The population-level growth-law response to translation inhibition predicts increased ribosome allocation as growth decreases (schematic arrow), whereas RpsB-YFP fluorescence decreases along the observed single-cell trajectories. PrpsL-CFP shows a distinct and more heterogeneous relationship with growth.

The two reporters responded in different ways. Following chloramphenicol exposure, growth rate rapidly decreased and the RpsB-YFP fusion signal progressively declined (Fig. 2b,c). Across tracked lineages, RpsB-YFP mean fluorescence was reduced during both intermediate (5 h) and prolonged (10 h) translation inhibition (Fig. 2d). PrpsL-CFP showed a more heterogeneous response. The PrpsL-CFP transcriptional reporter showed no significant difference at 5h and an overall reduction at 10h (Fig. 2d). However, its trajectory varied significantly between individual lineages (Fig. 2e). Chloramphenicol also increased cell length, width and projected area (Fig. SI2), so reporter signals were quantified using mean cellular fluorescence to account for treatment-induced changes in cell size. The decline in RpsB-YFP therefore cannot be explained simply by cellular enlargement. In contrast, total CFP fluorescence per cell increased after prolonged treatment, alongside cellular enlargement (Fig. SI2). <u>Together, these measurements show that the ribosomal protein fusion and transcriptional reporter follow distinct trajectories during translation inhibition</u>.

Examining the paired changes in both reporters within individual lineages revealed substantial heterogeneity in this response (Fig. 2e). Many lineages showed the predominant pattern of decreasing RpsB-YFP together with increasing PrpsL-CFP, but others exhibited weaker transcriptional induction, and a subset showed decreases in both reporters. Because both signals were measured simultaneously within the same tracked lineages, this divergence cannot be explained by growth-dependent changes in population composition. Instead, translation inhibition produced heterogeneous within-lineage trajectories in which the translational fusion and transcriptional reporter showed different fluorescence dynamics.

This behaviour was unexpected. Translation inhibition is well established to increase cellular investment in the ribosomal sector even as growth decreases, yet the ribosomal protein fusion moved in the opposite direction to that expected for a simple reporter of ribosome allocation. The simultaneous increase in ∼40% of the rpsL transcriptional reporter further showed that the decline in RpsB-YFP fluorescence did not reflect a corresponding suppression of rpsL promoter activity. Instead, translation inhibition separated the dynamics reported by the ribosomal protein fusion from those reported at the transcriptional level.

Despite this shift in its absolute response, the RpsB-YFP signal retained a relationship with cellular physiology. Across individual lineages, RpsB-YFP fluorescence remained positively associated with growth rate before treatment and during chloramphenicol exposure, whereas the relationship between the transcriptional reporter and growth was weaker and changed across the response (Fig. 2f). This positive reporter-growth relationship was also observed across different nutrient conditions and at an additional chloramphenicol concentration (Fig. SI3), indicating that it is not specific to the growth condition or antibiotic dose used in the main experiment. Thus, although the absolute trajectory of the ribosomal fusion reporter was inconsistent with its interpretation as a simple quantitative proxy for ribosome allocation during translation inhibition, the reporter still retained physiological information within this construct.

We next asked whether the decline in ribosomal fusion fluorescence was specific to RpsB-YFP or reflected a more general property of ribosomal protein reporters during translation inhibition. We therefore constructed a panel of translational fusion reporters using two ribosomal proteins, RpsB (S2) and RplI (L9), each fused to cyan, yellow or red variants of fluorescent proteins (Fig. 3a).

**Figure 3:**
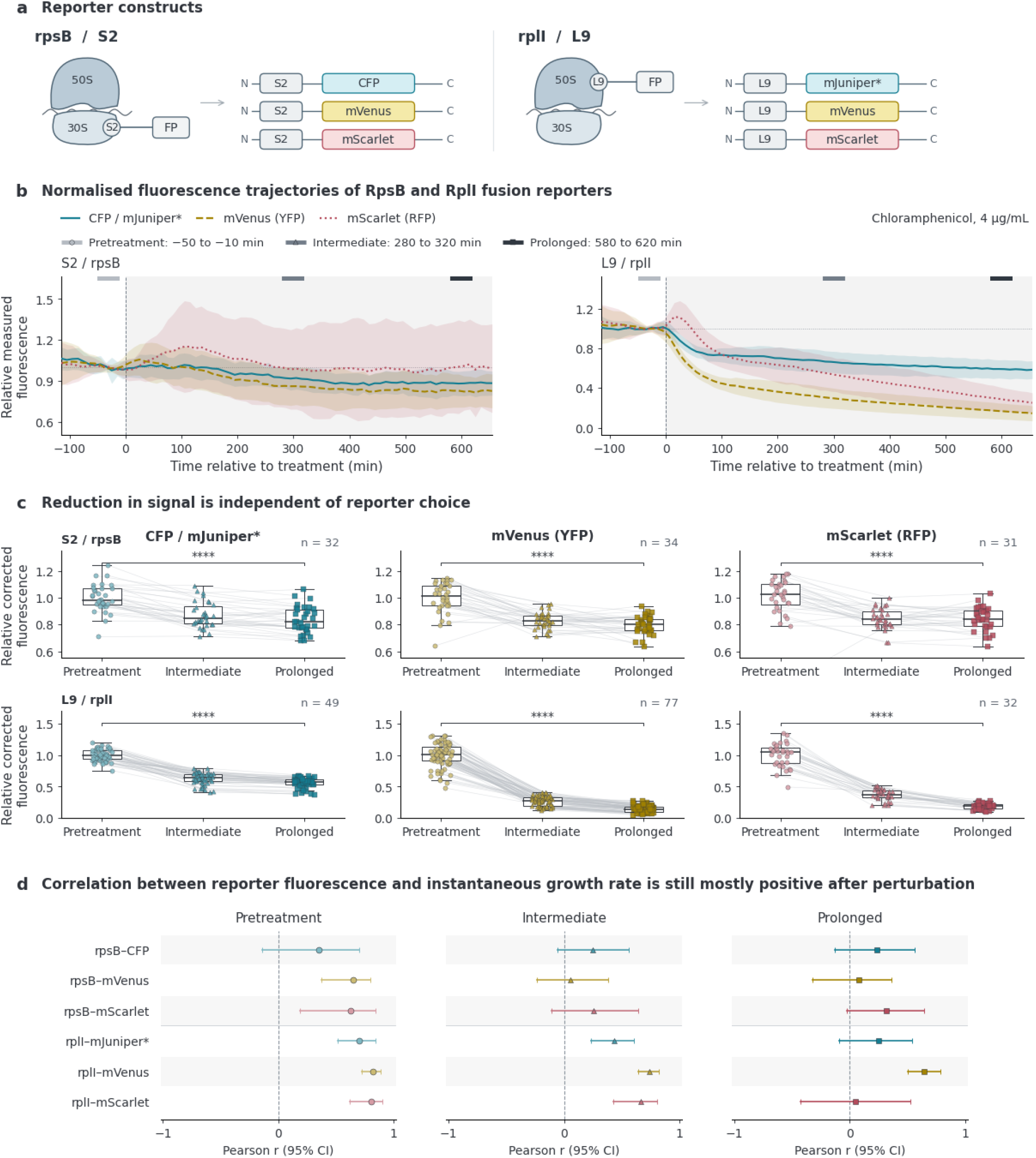
Ribosomal protein fusion fluorescence decreases across reporter architectures during translation inhibition. **a)** Schematic of the reporter panel comprising C-terminal fluorescent protein fusions to the ribosomal proteins RpsB (S2) and RplI (L9), each constructed with CFP, YFP and RFP variants. **b)** Normalised fluorescence trajectories of RpsB and RplI fusion reporters before and during chloramphenicol treatment (4 μg ml^−1^). Reporter fluorescence decreases following translation inhibition across both ribosomal proteins and all fluorescent proteins tested. Shaded bands show standard deviation. **c)** Change in reporter fluorescence between the pretreatment state and the intermediate and prolonged treatment periods for each construct, summarising the magnitude and consistency of the treatment-associated decline. **d)** Pearson correlation between reporter fluorescence and instantaneous growth rate across individual lineages before treatment and during intermediate and prolonged chloramphenicol exposure. Positive reporter-growth relationships are retained for some constructs but weaken during treatment for others, indicating that the physiological information carried by ribosomal fusion fluorescence depends on reporter architecture. Confidence intervals are shown in bars. Full reporter-growth scatter plots are shown in Fig. SI4.

Across this reporter panel, chloramphenicol treatment produced a consistent decrease in ribosomal fusion fluorescence (Fig. 3b). The effect was observed for both RpsB and RplI and across all three fluorescent proteins, although the magnitude of the decline varied between constructs, with the RplI/L9 reporters generally showing a larger decrease than the corresponding RpsB/S2 reporters (Fig. 3c). All strains showed comparable growth inhibition under the same treatment conditions (Fig. SI4), arguing against differences in antibiotic susceptibility as the explanation for these reporter-specific responses. <u>The reduction in fluorescence is therefore not restricted to a particular ribosomal protein or fluorophore.</u>

We then asked whether the relationship between reporter fluorescence and cellular physiology was similarly preserved across constructs. Before treatment, ribosomal fusion fluorescence was positively associated with instantaneous growth rate across the reporter panel. During chloramphenicol exposure, however, this relationship was retained to different extents across constructs (Fig. 3d; full scatter plots in Fig. SI4). Notably, the larger fluorescence decline of the RplI/L9 reporters did not correspond to a weaker physiological relationship; several L9 constructs retained a stronger positive correlation with growth than the corresponding RpsB/S2 reporters.

Thus, the magnitude of the treatment-induced shift in reporter fluorescence and the extent to which the signal preserves relative physiological information are separable properties of the reporter response. These within-treatment correlations should not be interpreted as the steady-state translation-limited growth law: the cells are undergoing dynamic adaptation, and the correlations describe relative variation among lineages within the same perturbation rather than the relationship between equilibrated states at different levels of translation inhibition.

Fluorescence measurements were corrected for growth-dependent dilution and fluorophore maturation using the measured growth rate and fluorophore-specific maturation time (see Methods). Nevertheless, residual reporter-specific measurement properties may contribute to the differences in correlation strength, including the larger autofluorescence contribution in the CFP channel and the slower maturation kinetics of RFP variants. Together, these results show that while <u>the decrease in ribosomal fusion fluorescence is robust across reporter architectures, the quantitative relationship between fluorescence and physiological state remains reporter-dependent.</u>

Finally, we asked whether the decrease in ribosomal fusion fluorescence was specific to chloramphenicol or extended to a second translation inhibitor. Tetracycline treatment similarly reduced RpsB-mVenus fluorescence across tracked lineages (Fig. SI5), despite its distinct mechanism of action. RpsB-mVenus fluorescence declined during tetracycline treatment in parallel with growth inhibition, reproducing the qualitative behaviour observed with chloramphenicol. Together with the reporter-panel analysis above, these results show that the decline in ribosomal protein fusion fluorescence is not restricted to a particular ribosomal protein, fluorophore or translation inhibitor. Thus, <u>even when population-composition effects are removed by lineage tracking, fluorescent ribosomal protein fusions can fail to reproduce the expected direction of ribosome regulation during translation inhibition.</u>

## Discussion

Quantifying physiological adaptation requires distinguishing changes that occur within individual cells from changes in the composition of the population through which those responses are observed. When the measured physiological variable is coupled to growth, differential growth progressively changes the representation of heterogeneous cellular trajectories within a population. Population-level measurements therefore reflect both regulation occurring within cells and growth-dependent reweighting of those cells over time, whereas lineage-resolved measurements follow the same initially sampled cells through the perturbation. These measurements answer different biological questions: the population average describes the state of the population present at a given time after differential growth has reshaped its composition, while the lineage average describes the mean trajectory followed by the initially sampled cells. For questions about regulation enacted within cells, the latter is the relevant quantity. Our model is deliberately simplified and is not intended to provide a quantitative description of the single-cell relationship between ribosome allocation and growth. Rather, it illustrates the general sampling effect that lineages with lower cumulative growth become progressively under-represented in an expanding population, causing population-level averages to diverge from the equally weighted trajectories of the initially sampled cells. Lineage tracking therefore provides a way to separate within-lineage regulatory dynamics from this growth-dependent population reweighting during translation inhibition.

However, our lineage-resolved experiments reveal a second and distinct limitation: even when the same cells are followed continuously, accurate physiological inference requires the reporter signal to remain quantitatively coupled to the biological variable it is intended to represent. Even within the same tracked lineages, fluorescent ribosomal protein fusions did not follow the expected direction of ribosome regulation during translation inhibition. Following chloramphenicol treatment, fluorescence from the RpsB/S2 fusion progressively decreased within individual lineages, despite the well-established increase in ribosomal allocation associated with translation inhibition. In contrast, a fraction of transcriptional reporters driven by ribosomal protein promoters increased during the same perturbation. Importantly, these opposing responses are observed within tracked lineages and therefore cannot be explained simply by differential growth changing population composition. The discrepancy is also not readily attributable to a particular fluorescent protein, ribosomal protein or antibiotic: similar behaviour is observed across multiple fluorescent proteins, with an independent ribosomal protein fusion, and during tetracycline treatment. Together, these observations indicate that the relationship between ribosomal protein fusion fluorescence and ribosomal physiology established under balanced growth does not remain invariant during translation inhibition.

Our experiments do not by themselves establish which molecular step produces this uncoupling. Fluorescence from a ribosomal protein fusion is the endpoint of several processes, including expression of the tagged protein, its incorporation into ribosomal particles, fluorescent-protein maturation and generation of the measurable fluorescent state. Perturbing translation could alter one or more of these relationships, such that fluorescence no longer scales with the underlying reporter abundance in the same manner as during balanced growth. The opposing response of the ribosomal transcriptional reporters is informative in this respect: the decline in fusion fluorescence cannot simply be interpreted as evidence that ribosomal gene expression has decreased. Thus, the mapping between fluorescence and the underlying ribosomal state should not be assumed to remain fixed during a perturbation that directly affects the machinery generating the reporter signal.

A useful way to formalise this problem is to distinguish the biological quantity of interest from the reporter-specific mapping that generates the measured fluorescence. We can represent the measured signal as *F*(*t*) = α(*t*)*X*(*t*), where *X*(*t*) is the underlying biological quantity and α(*t*) is the effective relationship between that quantity and fluorescence *F*(*t*). Note, this expression is intended as an effective representation of the reporter mapping rather than a kinetic model; in general, fluorescence may depend on the history of reporter synthesis and maturation as well as on its instantaneous abundance. Quantitative interpretation across conditions requires α(*t*) to remain sufficiently stable across the conditions being compared. Under balanced growth, this stability is often assumed implicitly. During translation inhibition, however, the perturbation directly alters the processes required to generate the fluorescent signal, including protein synthesis, growth-dependent dilution and fluorophore maturation. Our data therefore indicate that α(*t*) should not be assumed to remain invariant during the perturbation.

This distinction also helps explain why a reporter can remain physiologically informative without remaining quantitatively calibrated across conditions. In our measurements, some ribosomal protein fusions retain a positive relationship with growth within a given physiological state even though their absolute fluorescence trajectory is inconsistent with the expected direction of ribosome allocation. The strength of this relationship is itself reporter-dependent, weakening for some fluorophores during prolonged treatment. Thus, preserving relative information within a condition does not guarantee that the same fluorescence-to-physiology relationship can be extrapolated across a dynamic perturbation.

In a companion study, Zodage et al. address this question using biochemical measurements and an orthogonal fluorescence-LacZ reporter and find that fluorescence output can become uncoupled from reporter abundance during translation inhibition^10^ (see comparison in SI Table 1.). Their biochemical observations and our lineage-resolved measurements therefore identify complementary aspects of the same problem: the former establishes uncoupling between reporter abundance and fluorescence, while our measurements show how this discrepancy manifests dynamically within individual cells undergoing physiological adaptation.

These findings do not imply that fluorescent ribosome reporters are generally unreliable. Such reporters remain powerful for visualising ribosome organisation and relating ribosomal physiology to growth under defined conditions. The limitation is more specific: a fluorescence-to-physiology relationship established under one condition should not be assumed to remain quantitatively valid when the perturbation directly alters the processes that generate the reporter signal. This is especially important in dynamic experiments, where reporter synthesis, maturation, dilution and physiological adaptation occur on overlapping timescales. Under these conditions, fluorescence should be treated as a measured signal whose relationship to the biological variable of interest requires validation under the perturbation itself.

More broadly, our results identify two independent requirements for interpreting dynamic cellular measurements. First, the cells contributing to a measurement must represent the biological trajectories of interest; differential growth can otherwise cause population-level measurements to diverge from the trajectories followed by individual cells. Second, the reporter must continue to represent the physiological quantity being inferred as conditions change. Lineage tracking addresses the first problem but cannot by itself solve the second. Robust inference of regulatory dynamics therefore requires both longitudinal sampling of the relevant cellular trajectories and validation that the reporter-physiology relationship remains appropriate across the perturbation. This distinction should be relevant beyond ribosome regulation whenever the phenotype being measured is coupled to growth or the perturbation alters the machinery required to generate the reporter signal.

## Methods

### Agent-based simulation of growth-dependent population reweighting

The agent-based model was implemented in Python to examine how heterogeneous proliferation and differential growth affects population measurements of ribosome allocation. The population was initialised with five founder cells of equal cell size, each representing one lineage. Initial growth rates were assigned at evenly spaced intervals sampled from a uniform distribution with mean μ_*λ*_ (0. 8 *h*^−1 −1^) and standard deviation σ_*λ*_ (0. 18 *h*^−1^). Initial ribosome fractions were assigned using the linear relationship ϕ_*R, i*_ (0) = ϕ_*base*_ + *λ*_*i*_ (0)/κ_*t*_ where ϕ_*base*_ is the zero-growth intercept and κ_*t*_ is the growth-law coefficient. This initialisation provides a reproducible distribution of founder states with equal initial representation. It represents a defined population snapshot; differential proliferation can subsequently alter its composition even without treatment.

Translation inhibition was represented by a common residual growth factor q (= 0.6), giving the treatment-associated steady states: 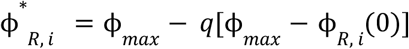. Treatment-associated ribosome fractions were assigned using a linear translation-inhibition relationship extrapolating to a shared maximum ribosome fraction at zero growth. These population-level growth-law relationships were used solely to define illustrative initial and endpoint states for each founder lineage; they were not assumed to constitute deterministic single-cell regulatory laws.^1^, each founder’s initial and treated states lie on an inhibition line extrapolating to the new shared intercept at zero growth (*λ*, ϕ_*R*_) = (0, ϕ_*max*_). With partial inhibition, treated growth rates and ribosome fractions remain heterogeneous. Applying these population-level growth-law relationships to individual founders was an illustrative modelling assumption. The relationships specified initial and final states, while the intervening adaptation trajectories were defined independently.

Three adaptation scenarios were compared using identical founder states and treatment-associated endpoints. In the instantaneous scenario (Fig 1b), cells adopted their treated physiological states at *t* = 0^+^, with their existing sizes retained. In the two gradual scenarios (Fig. S1a, b), either growth reduction preceded ribosome allocation, or ribosome allocation preceded growth reduction. Each response occupied a separate 3 h adaptation stage. Within a stage, adaptation was described by a smooth transition for both physiological trajectories:

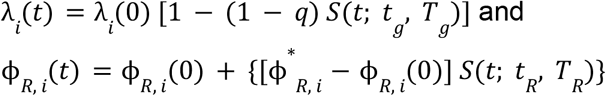

With a smooth cubic function which has zero slope at both stage boundaries between 0 and 1: *S*(*t*; *t*_0_, *T*) = 3*u*^2^ − 2*u*^2^, where 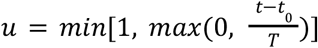. For growth-first adaptation, (*t*_*g*_, *t*_*R*_) = (0, 3) *h*; for ribosome-first adaptation, (*t*_*g*_, *t*_*R*_) = (3, 0) *h*; both have adaptation windows *T*_*g*_, *T*_*R*_ = 3 *h*. These schedules are illustrative limits on the single-cell response dynamic.

Cell size, V, used as a proxy for biomass, increased according to 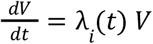. Division followed a deterministic adder rule: a cell divided symmetrically after increasing its size by a fixed increment, Δ, from its birth size^12^. All founders were initialised with birth size Δ and size 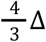 at the onset of treatment. Consequently, cells divided at size 2Δ, producing two daughters of size Δ. Daughter cells inherited their lineage’s physiological state and continued its adaptation schedule relative to the original treatment onset. No additional variability was introduced at division, and cell death, phenotypic switching, interactions between cells and resource limitation were excluded. We note that explicit cell division was included to visualise lineage expansion, but the population-reweighting effect is determined by differences in cumulative lineage growth and does not depend on specific division rule.

Under these assumptions, descendants within each founder lineage remained identical. Defining cumulative growth as 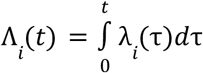, total lineage biomass and cell number were 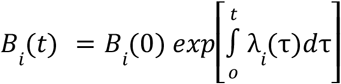 and 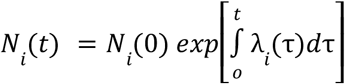. Growth integrals were evaluated analytically, and division events were determined from the adder thresholds. Accordingly, biomass increased continuously, whereas cell numbers changed discretely at division.

The population ribosome fraction plotted in the figures was calculated as the biomass-weighted mean, 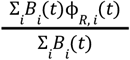, and compared with the equally weighted founder-lineage mean, 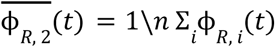. Cell-number-weighted means were also calculated by replacing *B*_*i*_ (*t*) by *N*_*i*_ (*t*). A homogeneous reference was initialised at the mean founder growth rate (0. 8 *h*^−1^) and ribosome fraction and subjected to the same adaptation law. This reference coincides with the equally weighted lineage mean. Simulations covered 12 h, with a 6h timepoint highlighted (Fig. 1d). Population composition could continue to change and converge to the fastest lineage after this time because differences in lineage growth persisted. All model parameters were chosen for illustration and were not fitted to experimental measurements. The full list is given in table 1.

### Strain construction

RpsB and RplI fusion strains were generated by λ-Red recombination in the background strains *E. coli* MG1655 CGSC 6300 (“SB1”) and *E. coli* MG1655 CGSC 7740 ΔmotA (“SB8”). Three different colour versions were created for each strain, with C-terminal fusions of either the CFP, SCFP3A^13^ or mJuniper^14^, the YFP, mVenus NB^13,15^, or the RFP, mScarlet-I3^16^. All versions included a (GGGGS)_3_ flexible linker between the ribosomal protein and the fluorescent protein. The genomic region surrounding *rpsB* or *rplI* was subsequently sequenced for confirmation for each version.

λ-Red recombination^17^ was performed using a modified version of the protocol described in^18^. Cultures of cells containing a gentamicin-resistance version of the pKD46 plasmid were prepared in LB broth (0.5 g/L NaCl) containing 10 μg/mL gentamicin and incubated overnight at 30 °C with shaking. The next day, the cultures were diluted 1:200 into fresh LB broth containing gentamicin and incubated at 30 °C with shaking until the OD600nm reached approximately 0.4, at which point L-arabinose was added to a final concentration of 0.3% (w/v) and the cultures were incubated at 37 °C with shaking for an additional 1 hour. The cultures were then transferred to ice, before being washed by four cycles of centrifugation and resuspension in ice-cold sterile distilled water. After the final centrifugation step, the liquid was discarded and the pelleted cells were resuspended in the residual liquid volume. The OD600nm of dilutions of the resuspended cells were measured and appropriate volumes of ice-cold sterile distilled water were added to bring the cells to a final density of OD600nm = 50.

Template plasmids were assembled by Golden Gate Assembly, comprising of the 500 bp homology region up to the end of *rpsB* or *rplI* (excluding the stop codon), a (GGGGS)_3_ flexible linker, the appropriate fluorescent protein, an FRT-flanked kanamycin resistance cassette or an FRT-flanked ampicillin resistance cassette, and the 500 bp homology region downstream of *rpsB* or *rplI*, within a pJUMP29-1A backbone^19^. This template was amplified using Q5 polymerase (NEB) and gel purified (NEB). 100 ng of each purified amplicon was added to 70 μL of the prepared pKD46-induced electrocompetent cells and transformed by electroporation. Cells were recovered in SOC media at 37 °C with shaking for 1-2 hours, then plated on selective LB agar plates containing 30 μg/mL kanamycin. The FRT-flanked kanamycin-resistance cassettes were excised from the strains using the plasmid pCP20, as described in^20^.

### Microfluidic device fabrication

Microfluidic devices were fabricated by standard polydimethylsiloxane (PDMS) soft lithography. PDMS base elastomer and curing agent (Sylgard 184, Dow) were mixed at a 10:1 (w/w) ratio, degassed under vacuum for 30 min, poured over the SU-8 master mould, and degassed again to remove residual air bubbles. The PDMS was cured at 95 °C for 1 h, peeled from the master, and cut into individual devices. Inlets and outlets were created using a 0.75-mm biopsy punch. PDMS replicas were cleaned by sonication in isopropanol for 30 min, dried with compressed air, baked at 95 °C for 30 min, and subsequently sonicated in deionised (DI) water for 30 min. Glass coverslips were cleaned by sonication in 1 M KOH for 20 min, rinsed, sonicated in DI water for a further 20 min, dried with compressed air, and baked at 95 °C for 30 min. PDMS devices were bonded feature-side down to glass coverslips following air-plasma treatment for 2 min at 35 W and a chamber pressure of 0.2–0.3 mbar using a Zepto plasma cleaner (Diener Electronic GmbH & Co. KG). Bonded devices were post-baked at 95 °C for 30 min and inspected under a stereomicroscope before use.

### Cell loading in the microfluidic device

Immediately before the experiments, the devices were passivated and cleaned by flowing M9 buffer + Pluronic F-108 (0.08% w/v) through the microfluidic lanes using a pair of gel-loading tips. Overnight cultures were grown in the presence of the same pluronic concentration to minimise cell aggregation and maintain uniform single-cell suspensions. For loading, the overnight cultures were passed through the flow lane at 5 ul min^-1^ for 1 minute, then fresh medium was flown to flush excess cells from the flow lane and to supply nutrients to cells retained in the trenches.

### Timelapse image acquisition

Timelapse microscopy was performed on a Nikon Eclipse Ti2 inverted microscope equipped with a Hamamatsu C14440-20UP camera. Images were acquired using a 40×/0.95NA phase-contrast (PC) objective and a 1.5x post-objective magnifier. The microfluidic device was fixed on the motorised stage inside a temperature-controlled incubator maintained at 37 °C, and continuously provided with media using a syringe pump connected via silicon tubing and 21-gauge needles. At the designated treatment time point, the inlet medium was then switched from fresh M9 medium to M9 supplemented with the required antibiotic concentration. Chloramphenicol working solutions were prepared by serial dilution from a 25 mg/mL stock solution made from chloramphenicol powder (Sigma-Aldrich), while tetracycline working solutions were prepared by serial dilution from a 50 mg/mL stock solution made from tetracycline powder (Thermo Fisher Scientific). Medium switching was achieved by a pinch valve system to prevent air bubbles. Where recovery was monitored, the treatment medium was subsequently replaced with antibiotic-free medium at the specified time point. Multiple fields of view (FOVs) were imaged automatically using the automated stage, with focus stability maintained by Nikon’s Perfect Focus System to compensate for thermal drift. Acquisition started 30 min after fresh M9 was introduced to stationary phase cells, and images were collected every 3 min. Fluorescence images were acquired every 3 min using a Lumencor Spectra X Light Engine at 50% excitation power and 100 ms exposure per channel. Excitation wavelengths were 438 nm (CFP), 510 nm (YFP) and 575 nm (RFP), with corresponding emission filters of 475/20 nm, 540/21 nm and 632/60 nm.

### Data processing for single-cell data analysis

Microscopy data were recorded in ND2 format and converted into per-FOV image stacks (lossless PNG stacks) using custom Python scripts. Time series were registered to compensate for any XY stage drift, and each FOV was cropped into individual trench movies using a custom-designed image-processing pipeline^9^. Single cell segmentation was performed on the trench movies using Cellpose-SAM^21^. Cell morphology was quantified using scikit-image’s regionprop package^22^. For each tracked cell, instantaneous growth rate was estimated from changes in cell length according to 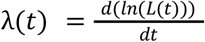, where *L*(*t*) is cell length at time *t*. In practice, 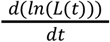 was calculated from the time-resolved cell-length trajectory smoothed by a first-order Savitzky-Golay filter with a 9-minute window between each division event. Prior to smoothing, abnormal spikes caused by segmentation error were flagged using a 1.3 fold-change threshold to detect when two large opposite changes surround one point while its neighbours remain consistent. Division events are flagged separately by a relaxed length drop ratio of 0.35 to 0.65. Growth-rate trajectories were aligned to the medium-switch time point to quantify changes before, during and after treatment. Fluorescence measurements were corrected for growth-dependent dilution and fluorophore maturation using a quasi-steady-state approximation, with a maturation dependent correction factor, such that *F*_*coss*_ = *F*_*measused*_ (1 + μτ), where μ is the instantaneous cellular growth rate and τ is the fluorophore maturation time. The maturation times were obtained from literature (SCFP3A: 7 mins^15^; mJuniper: 2 mins^14^; mVenus-NB: 4 mins^15^; YFP: 10 mins^5^; mScarlet-I3: 20 mins^14^; mCherry: 45 mins^14^).This correction assumes that reporter production and physiological state vary slowly relative to the maturation timescale.

### Statistical analysis

Statistical analyses were performed in Python using custom scripts. Unless otherwise stated, individual mother-cell lineages were the units of analysis for treatment state comparison and the units of resampling for bootstrap analysis. Repeated measurements obtained from successive imaging frames of the same lineage were not treated as independent observations.

For comparisons between physiological states, measurements were averaged within predefined time windows for each lineage. These corresponded to the pre-treatment period (−50 to −10 min relative to antibiotic addition), an intermediate-treatment period (280–320 min), and a prolonged-treatment period (580–620 min). Only lineages for which measurements were available in all time windows included in a paired comparison were analysed. A window mean was retained only when it contained at least five valid measurements. Differences between states were assessed using two-sided Wilcoxon signed-rank tests. For the dual-reporter experiment, tests were applied to paired differences in corrected fluorescence means. For comparisons across the six different ribosomal protein fusion reporters, treatment responses were expressed as *log*_2_ fold changes relative to the pretreatment fluorescence of the same lineage.

Associations between reporter fluorescence and cellular growth rate were quantified using Pearson’s correlation coefficient (*r*). For comparisons at defined physiological states, reporter fluorescence and growth rate were independently averaged within the corresponding time window for each lineage, such that each lineage contributed a single paired observation to the correlation analysis. Linear regression lines shown in scatter plots were included to visualise the direction and magnitude of these associations.

For time-resolved correlation analyses, reporter fluorescence and growth rate were averaged within a sliding window of 15 mins centred at each time point, and Pearson’s *r* was calculated across mother-cell lineages. All eligible measurements belonging to each sampled lineage were retained together, preserving their within-lineage dependence. Confidence intervals were estimated by non-parametric bootstrap resampling of complete mother-cell lineages with replacement (2000 bootstrap replicates).

Resampling was performed at the lineage rather than frame level to preserve the temporal dependence of repeated measurements obtained from the same cell. The 2.5th and 97.5th percentiles of the bootstrap distribution were used to define 95% confidence intervals.

For analyses comparing treatment-induced responses within the same cell, changes in reporter fluorescence were calculated relative to the corresponding pretreatment value for each lineage. Δ*F* = (*F*_*tseat*_ − *F*_*pse*_)/*F*_*pse*_. Associations between changes in different reporters measured within the same lineage were quantified using Pearson’s *r*.

Where multiple related hypotheses were tested, *P* values were adjusted using the Holm method within each predefined family of comparisons. Exact sample sizes and adjusted *P* values are reported in the corresponding figures or figure legends. All statistical tests were two-sided. Time-course summaries show the mean across mother-cell lineages, with variability represented by the standard deviation unless otherwise indicated. For estimated statistics such as time-resolved correlation coefficients, uncertainty is instead reported using bootstrap 95% confidence intervals. Throughout the manuscript, *n* denotes the number of independent mother-cell lineages unless explicitly stated otherwise.

## Acknowledgements

This work was supported by a University Startup Award for Lectureship in Synthetic Biology (NKXY ISSF3/46), an EPSRC New Investigator Award (EP/W032813/1), and an ERC Consolidator Award (Grant Number: G526637).

## Supplementary information

**Figure SI1:**
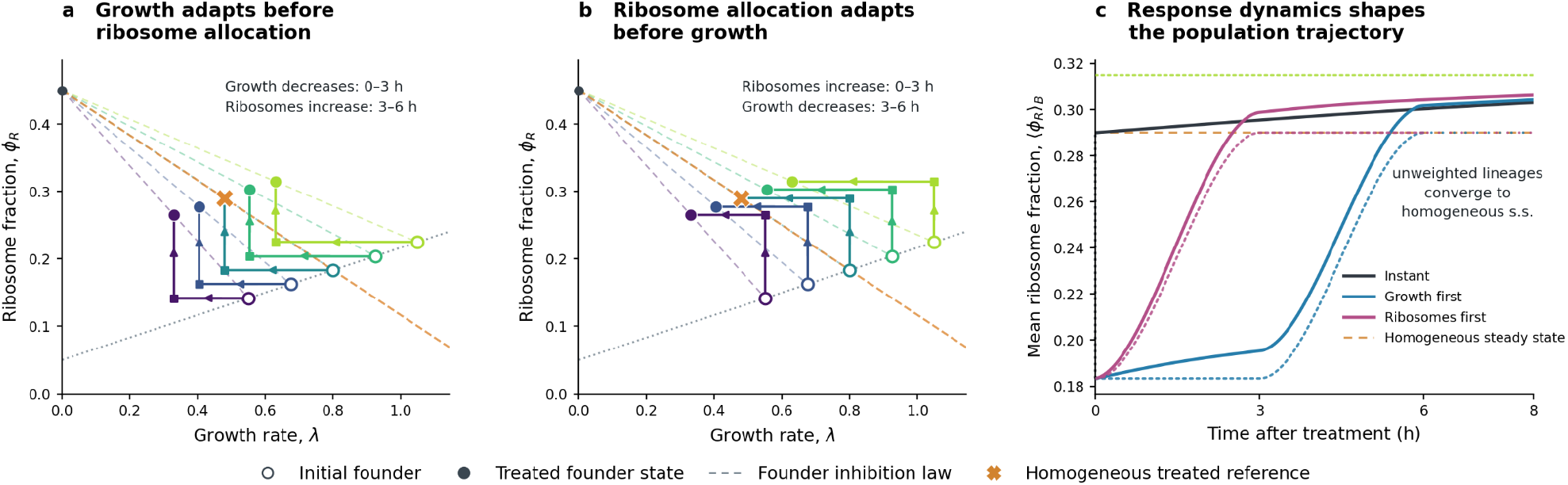
The divergence between population- and lineage-level measurements persists across different orders of physiological adaptation. **a,b)** Illustrative lineage trajectories when growth rate responds before ribosome allocation (a) or ribosome allocation responds before growth rate (b). Founder lineages begin from the same heterogeneous pretreatment states and approach the same treatment-associated endpoints as in Fig. 1, but traverse different paths through physiological state space depending on the ordering of the two responses. **c)** Corresponding population-average ribosome fraction over time for the two response scenarios shown in (a) and (b). Differential growth reweights the contribution of individual cellular trajectories, causing population-level measurements to depend on both the regulatory trajectories followed within cells and their associated growth histories. The lineage-resolved mean instead gives equal weight to the initially sampled cellular trajectories (dotted line). Thus, population and lineage measurements describe distinct quantities: the state of the evolving population and the mean regulatory strategy of individual cells, respectively. Model parameters are given in table 1.

**Figure SI2.**
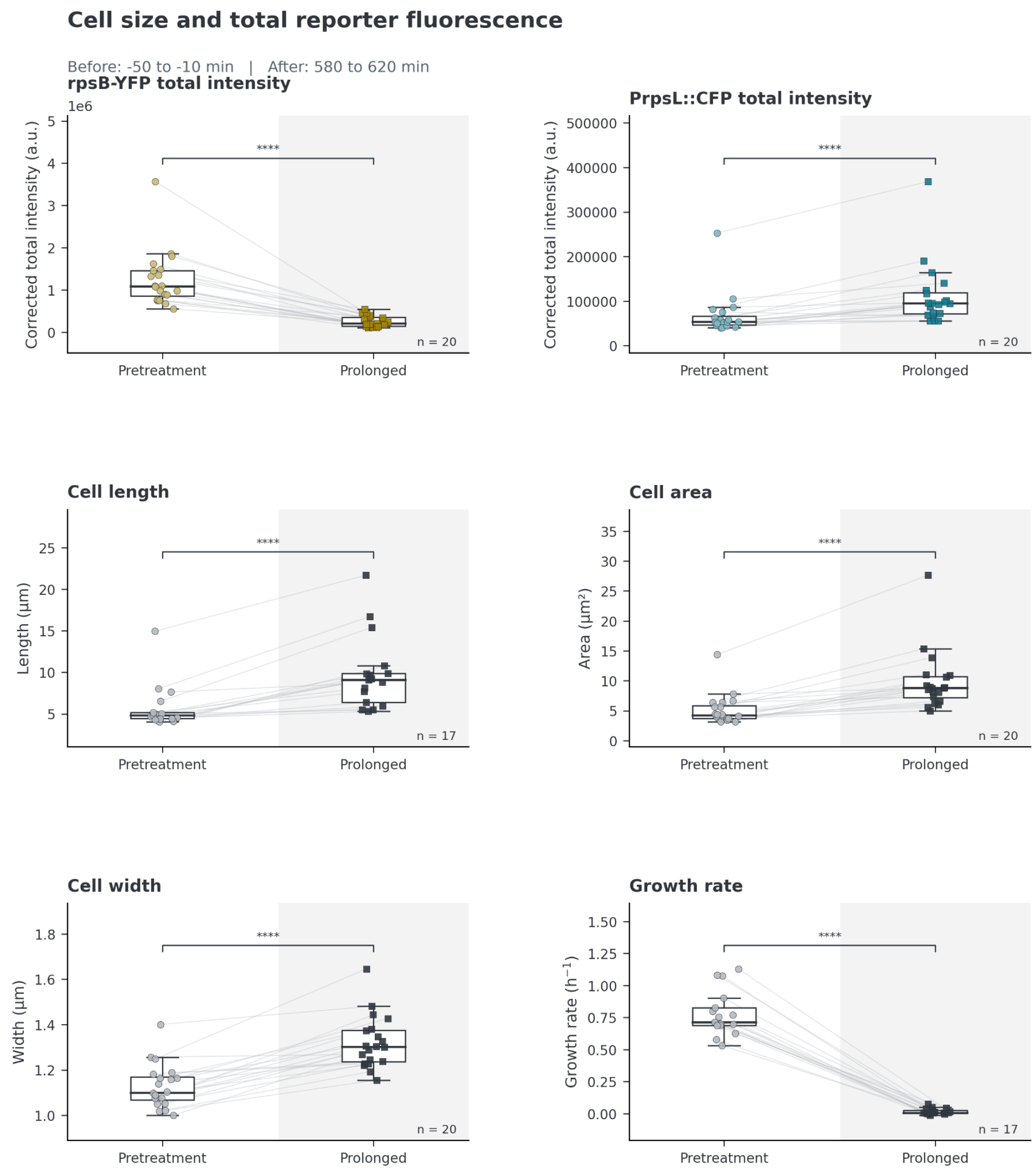
The decrease in RpsB-YFP fluorescence during chloramphenicol treatment is not explained by treatment induced changes in cell size. Cell length, width and projected cell area increase during translation inhibition across the reporter strains. These changes are important for quantitative interpretation of fluorescence measurements because increases in cell size can alter the relationship between total fluorescence and intracellular reporter concentration. Reporter analyses in the main text therefore use mean cellular fluorescence to account for treatment-induced changes in cell geometry. Growth-rate responses are shown alongside morphology to confirm the expected physiological response to chloramphenicol. Drop in total intensity of the RpsB fusion reporter confirms that the drop in mean RpsB-YFP signal is not related to cell size increase during treatment.

**Figure SI3:**
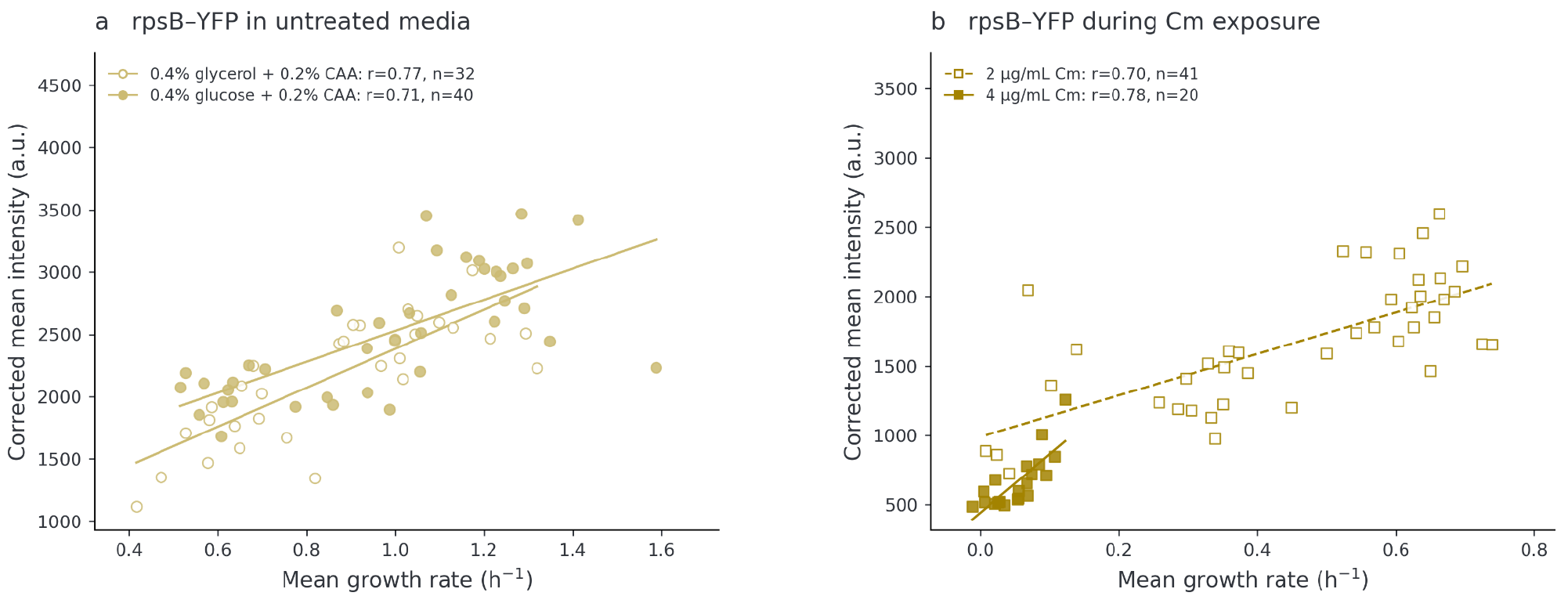
The positive association between RpsB-YFP fluorescence and cellular growth is preserved across nutrient conditions and chloramphenicol concentrations. **a)** Relationship between RpsB-YFP mean fluorescence and single-cell growth rate across cell-lineages growing in different nutrient conditions. The line indicates the linear regression. **b)** Relationship between RpsB-YFP mean fluorescence and single-cell growth rate during treatment with two different chloramphenicol concentrations. Pearson correlation coefficients (r) are shown for each case. In each of these scenarios, RpsB-YFP mean fluorescence remains positively associated with cellular growth.

**Figure SI4:**
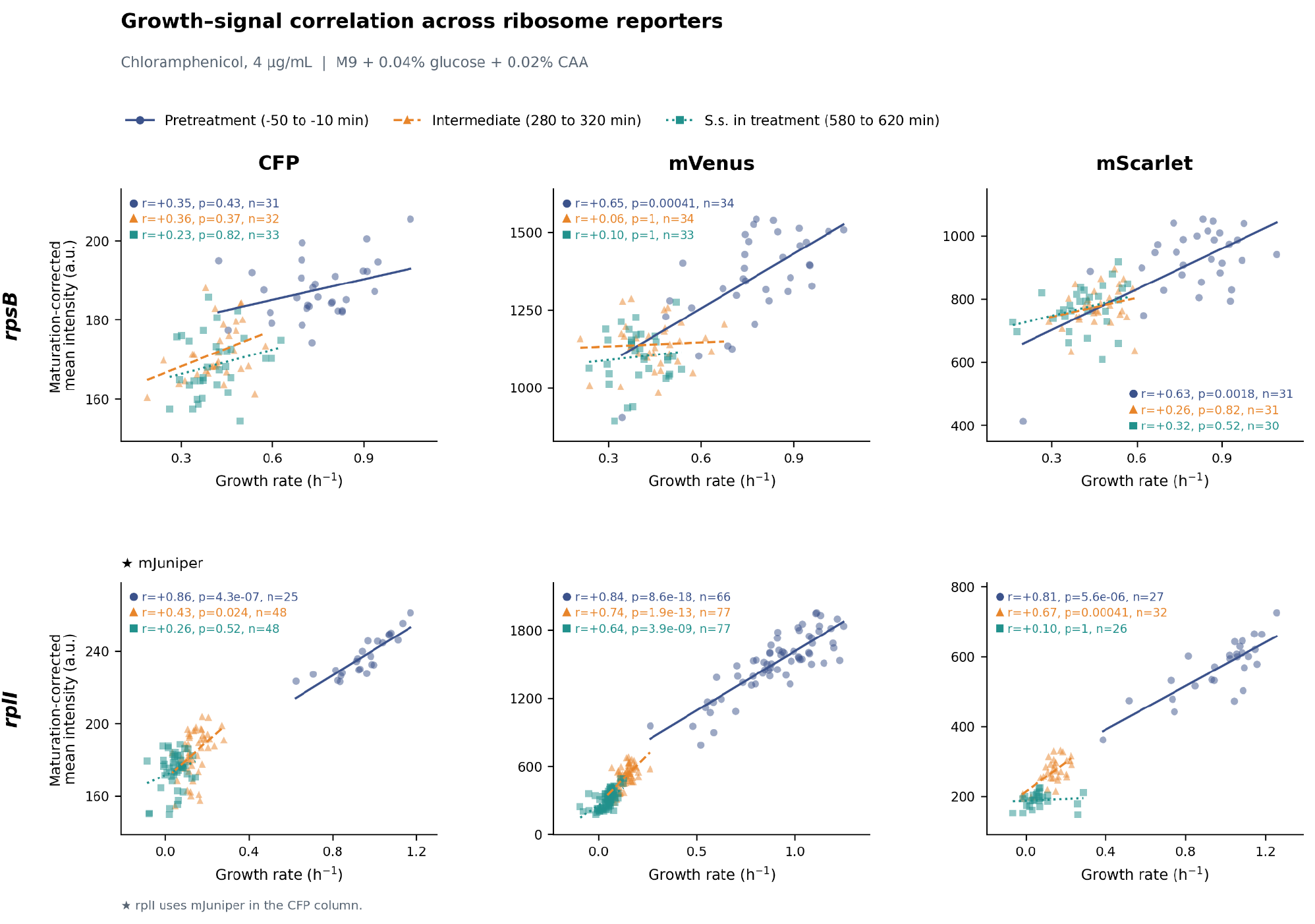
Reporter-growth relationships across ribosomal protein fusion variants during chloramphenicol treatment. Scatter plots showing the relationships between instantaneous growth rate and corrected mean fluorescence intensity for RpsB(S2) and RplI (L9) fusion reporters carrying CFP (mJuniper), YFP (mVenusNB) or RFP (mScarletI3) variants.Measurements are shown for the pretreatment period (−50 to −10 min), intermediate treatment (280–320 min) and prolonged treatment (580–620 min) during exposure to chloramphenicol (4 μg ml^-1^. Each point represents the mean fluorescence and growth rate of one mother-cell lineage within the corresponding time window. Pearson correlation (*r*) coefficient and associated *P* values are shown. Positive reporter-growth relationships are evident for several constructs before treatment and are retained to different extents during translation inhibition, with weaker relationships emerging for some reporters at later times. These data underlie the correlation coefficients and confidence intervals shown in Fig. 3d.

**Figure SI5:**
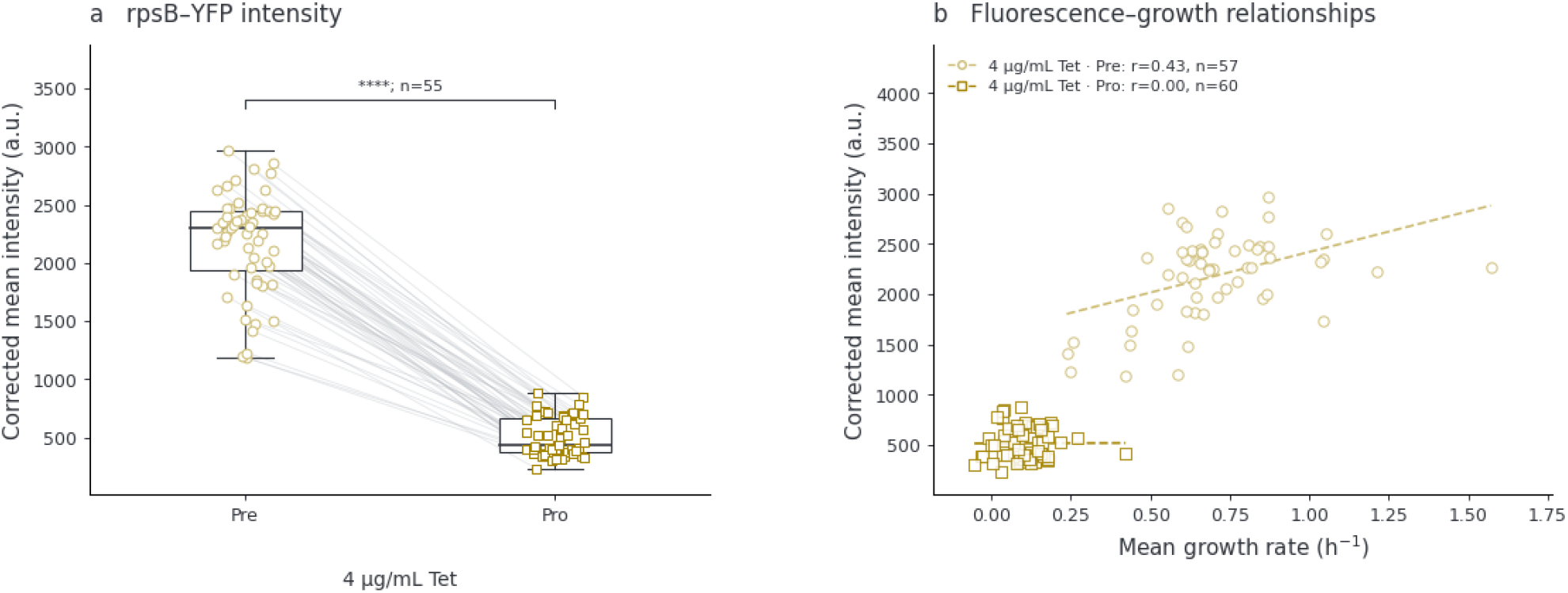
Tetracycline reproduces the decline in ribosomal fusion fluorescence observed during chloramphenicol treatment. **a)** Corrected mean fluorescence intensity of the RpsB-YFP ribosomal fusion reporter before and during treatment with tetracycline (4 μg ml^−1^). Measurements from the same tracked lineages are connected by grey lines. Tetracycline treatment produces a substantial reduction in ribosomal fusion fluorescence, reproducing the qualitative response observed during chloramphenicol exposure. **b)** Relationship between corrected RpsB-YFP fluorescence and instantaneous growth rate before and during tetracycline treatment. Each point represents an individual lineage and lines indicate linear fits; Pearson correlation coefficients and associated P values are shown. Tetracycline therefore reproduces the qualitative decrease in ribosomal protein fusion fluorescence observed during chloramphenicol treatment.

**Table S1.** Summary of experiments in the present study and Zodage et al.^10^.

|  | Present Study | Companion Study (Zodage et al.) |
| --- | --- | --- |
| **E. coli Strains** | RpsB-SCFP3A<br>RpsB-mVenus-NB<br>RpsB-mScarlet-I3<br>RplI-mJuniper*<br>RplI-mVenus-NB<br>RplI-mScarlet-I3<br>RpsB-YFP<br>PrpsL-SCFP3A<br>PrpsL-mVenus-NB<br>PrpsL-mCherry | RpsB-mChartreuse (SJ2524)<br>*rpsB::mChartreuse* (SJ2570)<br>*rpsB::lacZ* (SJ2571)<br>*rpsB::mChartreuse-lacZ* (SJ2531)<br>PrrnB::mChartreuse (SJ2560)<br>PrrnB::lacZ (SJ2571)<br>PrrnB::mChartreuse-lacZ (SJ2574)<br>PrrnE::mChartreuse (SJ2561) |
| **B. subtilis Strains** | — | RpsB-mChartreuse (JDW94) |
| **Media** | 2 growth media | 5 growth media |
| **Antibiotics** | Chloramphenicol, Tetracycline | Chloramphenicol |
| **Antibiotic Concentrations** | 1 to 2 concentrations per antibiotic | 4 to 5 concentrations per strain |
| **Measured parameters** | Growth rate<br>Fluorescence intensity (microscopy)<br>Cell morphology parameters | Growth rate<br>Fluorescence intensity (microscopy)<br>RNA-to-protein ratio<br>LacZ activity |
| **Replicates** | Parallel quantifications in the Mother Machine (N>20) | 1-to-5 replicates |

